# Epigenomic profiling of active regulatory elements by enrichment of unmodified CpG dinucleotides

**DOI:** 10.1101/2024.02.16.575381

**Authors:** Luca Tosti, Calum Mould, Imogen Gatehouse, Anyela Camargo, Anthony C. Smith, Krystian Ubych, Peter W. Laird, Jack Kennefick, Robert K. Neely

**Affiliations:** Tagomics Ltd, Cori Building, Granta Park, Cambridge, CB21 6GQ, UK; Van Andel Institute, 333 Bostwick Ave. NE, Grand Rapids, MI 49503, USA; The University of Birmingham, School of Chemistry, Edgbaston, Birmingham, B15 2TT, UK

## Abstract

Current approaches for the study of DNA methylation and other modified bases focus on the modified fraction of the genome and are particularly well-suited to the detection of DNA *hyper*methylation. However, the study of *hypo*methylation (loss of DNA methylation), which is typically associated with markers of active chromatin, has been largely overlooked, in part, due to the lack of a suitable methodology. We present an enrichment-based and bisulfite-free approach for epigenomic profiling named “Active-Seq” (**A**zide **C**lick **T**agging for **I**n **V**itro **E**pigenomic **seq**uencing) that achieves genome-wide profiling of DNA, by enriching for non-modified CpG sites using a mutated methyltransferase enzyme. We show that the genomic regions enriched by Active-Seq overlap with promoters, enhancers and partially methylated domains, all of which have had their methylation status linked to the development and progression of diseases. Active-Seq is a fast epigenetic profiling platform with a simple and streamlined workflow, performed in tandem with sequencing library preparation. The enzymatic chemistry is non-damaging toward the DNA which is critical for working with low concentration DNA input and will facilitate the future development of multiomics assays. We demonstrate robust and reproducible performance of Active-Seq using low DNA input in cell lines, liquid biopsies (cell-free DNA, cfDNA) and formalin-fixed paraffin embedded (FFPE) tissue.

## Introduction

Epigenetic modifications of DNA, such as methylation and hydroxymethylation of cytosine, play a pivotal role in gene regulation^1^. Indeed, methylation of DNA is critical in embryogenesis, early development and it is known to change predictably in correlation with biological ageing of an organism^2–4^. On the other hand, aberrant methylation of DNA is an important driver of tumorigenesis as the epigenetic alterations of gene regulatory regions play a key role in many diseases^5–8^.

The established link between DNA methylation and disease, as well as the relative stability of this covalent DNA modification, makes it an ideal candidate biomarker. However, the measurement of DNA methylation levels remains challenging. Large-scale, genome-wide, comparative studies of the human DNA methylome have been largely carried out using array-based platforms, with the latest array covering 935,00 sites, i.e. around 3% of the 28M CpG sites of the human genome^9^. The focus on a small fraction of the DNA methylome limits the discovery of new methylation-based biomarkers and the study of large-scale regions of contiguous (un)methylation. Moreover, genome-wide, comparative technologies (using bisulfite conversion, EM-Seq^10^ or TAPS^11^) can be prohibitively expensive at the population scale. Despite this, recent work by Loyfer and colleagues demonstrates the value of a genome-wide approach (using whole genome bisulfite sequencing, WGBS) applied to 77 primary cell types in healthy human cells to discover striking differences in the methylation patterns of healthy cells, particularly in unmethylated regions of the genome^12^. These novel epigenetic markers are blocks of unmethylated DNA, specific to different cell types and their progeny and are strongly correlated with markers of active chromatin (active enhancers, H3K27ac and H3K4me1^12^). The recent discovery of these DNA methylation-based biomarkers after over twenty-five years of research effort in this space, highlights the need to increase the spectrum of technologies that can be applied to facilitate the study of DNA methylation, and fully resolve its complexity.

Extended global hypomethylation of the genome was the first epigenetic process to be linked with cancer development^13,14^. Since, it has been shown that this process is driven in regions of the genome that replicate late in the cell cycle and the concomitant rapid cell division that occurs in tumours^15–19^. This feature of the cancer genome has been linked to the early developmental stages of cancer through studies of methylation at LINE-1 (as well as other repetitive genomic) elements, that act as a proxy for the ‘genome-wide’ methylation level^20–24^. The emerging picture reveals that DNA hypomethylation, or the lack of DNA methylation, is an activating marker that could be a critical driver and/or indicator of cell biology. However, this is at odds with the current techniques for profiling DNA methylation, which ubiquitously focus on the detection of methylated bases.

The early detection and diagnosis of cancer is moving towards processing of samples from liquid biopsy, and particularly from blood^25,26^. Cell-free DNA (cfDNA) is present in the blood plasma of both healthy and diseased individuals and can be composed of DNA shed from healthy cells, as well as circulating tumour DNA (ctDNA) from dying tumour cells^27^. In principle, analysis of ctDNA will enable personalised cancer treatment, tailored to the (epi)genetic makeup of the tumour. The challenges of liquid biopsy are twofold i) the low concentration of cfDNA in the blood (typically nanogram quantities per millilitre of plasma), and ii) the small proportion of cfDNA that is tumour derived^28–30^. Hence, there is a pressing need for the development of an epigenetic technology compatible with minute DNA input quantities (non-damaging, sensitive and efficient), that can be applied genome-wide and across the spectrum of DNA methylation markers of disease (hyper- and hypo-methylated sites) for cost-effective diagnosis that can be deployed on a population scale. Recent developments in enrichment-based technologies have begun to address this challenge^5,31^ but both antibodies for 5-methylcytosine and MBD proteins are known to display preferential binding towards heavily methylated regions of the genome^32^.

To address the limitations of current epigenomic technologies, we present Active-Seq, an enzymatic technology for the enrichment of unmethylated DNA molecules. Active-Seq builds upon pioneering work from by Kriukienė et al.^33^ and employs an engineered bacterial DNA methyltransferase enzyme to specifically target and tag unmethylated CpG sites across the whole genome. We show that Active-Seq results in robust enrichment of unmethylated CpG-containing DNA. The workflow is strikingly simple and can be integrated readily with standard library preparation in a single pot reaction. We show that Active-Seq enables robust epigenomic profiling using sub-10 ng amounts of input DNA. The inherent simplicity of the approach means that Active-Seq is robust across laboratory operators and sequencing runs and can be used to generate genome-wide epigenetic profiles, with no assumed knowledge of the sample. An additional benefit of the approach is that enzymatic tagging does not damage DNA, meaning that mutational signature is preserved in the sample, as is the fragmentomic information, i.e. features such as fragment size, end motif sequences, etc.^34,35^. Active-Seq has the potential to serve as the foundation for a streamlined, multiomic and clinically deployable diagnostic tool, particularly where only low DNA input amounts are available such as in liquid biopsy.

## Results

Active-Seq approach is described schematically in Figure 1A. In short, we employ a bacterial DNA methyltransferase enzyme (M.MpeI) to tag non-modified CpG sites. We substitute the enzyme’s natural cofactor, *S*-adenosyl-L-methionine (AdoMet), with an unnatural cofactor analogue (azide tagging compound), that enables DNA transalkylation with an azide-terminated functional group. Incubation of the methylase, DNA and cofactor for one-hour results in near-complete modification of the target DNA (Supplementary Figure S1). These tags can be further modified (e.g. biotinylated) using highly efficient, bio-orthogonal click chemistry to enable enrichment of non-modified DNA using streptavidin-coated beads (Figure 1A). Note that ‘non-modified’ refers to all genomic DNA fragments containing at least one CpG dinucleotide that is not modified (methylated, hydroxymethylated, carboxylated, formylated) at the C5-position.

**Figure 1.**
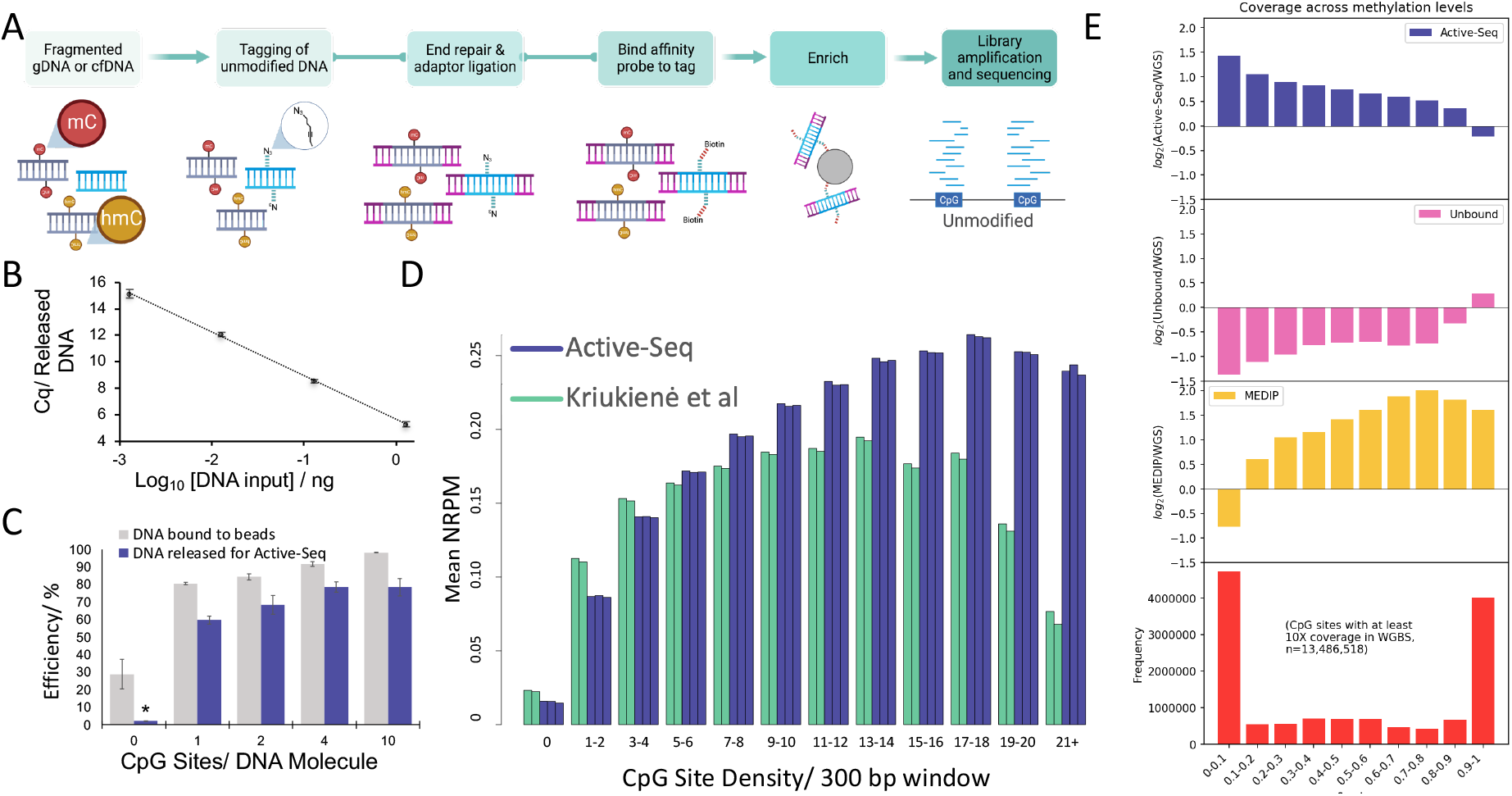
Active-Seq workflow and comparison with other epigenomics technologies. **(A)** Schematic showing the one-pot Active-Seq workflow. Purified DNA (cfDNA or fragmented gDNA) is tagged using a CpG-targeting methyltransferase (M.MpeI) and a synthetic cofactor analogue. The enzyme catalyses DNA alkylation with azide-terminated tags exclusively at unmodified CpG sites. Tagged DNA molecules are subsequently subject to standard library end-repair and adapter ligation, followed by biotinylation and isolation using streptavidin-coated magnetic beads. Tagged (unmodified CpG sites, Active-Seq) and untagged (5mCpG and other modified CpG sites, unbound fraction) DNA can be separately amplified and sequenced. **(B)** Plot showing binding and recovery efficiency for DNA oligos (150bp fragment containing 10 CpG sites) against a background of DNA containing no CpG sites. The plot shows qPCR threshold cycle versus target DNA concentration. **(C)** Binding efficiencies (grey) and recovery efficiencies (blue) of target DNA oligos as a function of target DNA CpG site density. * Indicates that DNA concentration of the unbound material was below the limit of detection. **(D)** Normalised Reads Per Million (NRPM) of genomics DNA as a function of CpG density in a given 300 bp window (human genome) for Active-Seq (blue) and for the method reported by Kriukienė *et al*. (green). **(E)** Log_2_ ratio of specific coverage vs WGS coverage for Active-Seq (blue), the Active-Seq unbound (‘methylated’) fraction (pink), MeDIP-Seq (yellow) as a function of CpG site β-value, as determined by WGBS (red) in NA12878 cell line DNA.

We have developed a single-tube approach for tagging, enrichment and DNA library preparation workflow, with the overarching aim of minimising handling and purification steps. The streamlined Active-Seq workflow, its overall efficiency and unbiased coverage of the whole methylome set it apart from other epigenetic approaches. The approach transforms the previously published approach for unmethylome profiling^33^ by minimising the loss of densely modified DNA from the sample, thereby retaining uniform representation of DNA fragments across the genome.

### Methyltransferase-directed enrichment of DNA is selective and efficient

We sought to establish the performance of Active-Seq as a function of DNA concentration and CpG density. Both facets of the experiment are critical for the application of the approach in challenging samples, such as in liquid biopsy. Our focus during development of the method was on the deployment of water-soluble chemistries for DNA enrichment and on minimising the number of DNA purification steps. The latter invariably requires condensation of the DNA and can lead to loss of DNA molecules which are densely tagged, biologically relevant, but sparingly soluble. To this end, we initially validated the performance of the enzymatic enrichment across a range of DNA concentrations and on DNA molecules containing a known range of CpG site densities. We spiked between 1.25 ng and 1.25 pg (equivalent to approximately 200 copies and 0.2 copies of a diploid, human genome, respectively) of these target DNA molecules (153 bp, containing 10 unmodified CpG sites) into a background of 24 ng of non-target DNA (142 bp containing no CpG sites). We tagged the target DNA and enriched this fraction using streptavidin-coated beads. The overall efficiency of the enrichment was determined by qPCR, using primers specific to the target DNA molecules (Figure 1B). We found DNA binding and recovery efficiencies in excess of 80% for all the samples, clearly demonstrating the compatibility of the approach for enrichment of DNA at input levels consistent with cell-free- or single-cell DNA analysis. Notably, we found that tagged DNA is directly compatible with PCR and can be amplified using a standard polymerase enzyme, with an efficiency that is on a par with that observed for unmodified DNA.

In a second validation experiment, we examined the binding and recovery efficiency of DNA at a range of CpG densities, from approximately 1 site/150 bp to 10 sites/150 bp, the latter of which is consistent with the CpG density found in CG islands. The initial step of binding of biotinylated DNA to streptavidin-coated beads shows a modest dependence on the number of CpG sites available for tagging on a DNA molecule, at low CpG site concentrations of less than 4/150 bp (Figure 1C). We also noted some non-specific binding of non-target DNA to the magnetic beads used for enrichment, though less than 10% of the non-specifically bound DNA is recovered from the beads. Recovery efficiencies show no detectable dependence on the CpG site density and overall binding and recovery efficiency of DNA is between 60% and 80%, across the range of tested CpG densities (Figure 1C). In contrast to the work by Kriukienė *et al*,^33^ we achieved highly efficient binding and recovery of all unmodified DNA molecules, regardless of the number of unmethylated CpG sites they carry. This enables the recovery of unmethylated CpG-dense promoter CpG islands, which may be lost from the profile using the earlier approach. We attribute this to the streamlined workflow for Active-Seq that maximises DNA solubility and minimises the exposure of highly modified DNA to conditions that might cause its precipitation (e.g. biotin is sparingly soluble in aqueous solution). Under the saturating conditions employed for our tagging reaction, we observe three-to-four times the binding and recovery efficiency of this previous work (Figure 1D). Active-Seq is consistently performant under saturating enzymatic labelling conditions, as we will show for genome-wide epigenomic profiling in more complex genomic samples below.

### An integrated workflow for epigenomic sequencing

Having demonstrated the performance of the biochemical approach on model DNA oligos, we sought to prove the utility of our enzymatic enrichment on genomic DNA and combined this with sequencing library preparation in a single pot workflow (Active-Seq). Genomic DNA from NA12878 cells was fragmented and we generated DNA libraries for the enriched (unmodified) and unbound (modified) fractions of the genome using Active-Seq. For the enriched, unmethylated DNA fraction, saturation analysis shows that signal reaches 90% saturation at approximately 70M reads (approximately 15Gbp of sequencing per sample, Supplementary Figure S2). Active-Seq focusses on enriching the 20-30% of the CpG sites in the genome that are unmethylated, leading to a dramatic reduction of the required sequencing depth compared to genome-wide approaches using base conversion. Processed reads in Active-Seq also show low levels of read duplicates (∼4.2%) that are significantly below the levels observed in bisulfite sequencing (about 30-40%) and EM-Seq (∼10%)^10^. Therefore, Active-Seq facilitates genome-wide epigenetic profiling with approximately 1/10^th^ of the sequencing depth required for base conversion technologies.

Successful enrichment at CpG sites, as a function of their methylation status, was assessed by further comparison of the sequencing results from the enriched (non-modified) and unbound (modified) fractions of the genome with β-values (average methylation levels) of CpG sites derived from whole-genome bisulfite sequencing (Figure 1E). Using CpG sites that have at least 10x coverage in the WGBS dataset, we observe significant enrichment (∼3x relative to whole genome sequencing) in Active-Seq at sites, with β-values of less than 0.1. This contrasts with the unbound fraction of the sample, where the WGS coverage is around a factor of three higher for these unmethylated CpG sites. Enrichment in Active-Seq falls gradually with increasing methylation levels until we see higher signal in the WGS experiment for highly methylated CpG sites, with β-values greater than 0.9. Hence, we can conclude that the Active-Seq profile shows high enrichment efficiencies for unmodified CpG sites, near uniform enrichment efficiency across a range of CpG densities, and no significant off-target enrichment of DNA. These initial validation experiments confirm the robustness of the enzymatic chemistry across a wide range of DNA substrate complexities.

### Active-Seq can be applied on challenging clinical biosamples

We sought to prove the feasibility of the approach using DNA derived from clinical blood (plasma) samples and from formalin-fixed and paraffin embedded (FFPE) tissue samples. We have shown that Active-Seq is performant with input DNA orders of magnitude less than one nanogram and, combined with the genome-wide aspect of the approach, anticipated that the analysis of ctDNA would be readily achievable. To confirm this, we processed a cfDNA sample from a healthy patient and one from a patient diagnosed with stage 1 lung cancer (non-small cell lung cancer). We performed Active-Seq in duplicate (separate preparations and sequencing runs). Both input DNA samples and the output of the enriched library maintain the fragment size distribution that is characteristic of nucleosomal cell-free DNA (Supplementary Figure S3). Duplication rates for the sequencing data were 9.2% and 9% for the larger sequencing run, with coverage of 100M reads for both samples. This represents a significant improvement on duplication rates typical for approaches using base conversion, which can reach 30-40% in studies of cfDNA^36^. The background, defined as the percentage of reads lacking a CpG site in the dataset, was 4.5% and 3.6% for the healthy and cancer samples, respectively. Cell-free Active-Seq profiles show remarkable consistency, despite the small quantity of input DNA (Supplementary Figure S4), highlighting the potential of the approach for application in liquid biopsy. We examine the possibility of deriving meaningful biological signal from this sample type below.

FFPE treatment typically leads to extensive damage (depurination, depyrimidation and deamination) of the genome. We were keen to establish the impact of this on Active-Seq and our ability to generate a meaningful epigenomic profile from DNA extracted from FFPE tissue. We generated genome-wide Active-Seq profiles for three FFPE embedded samples in triplicate, sourced from the Wales Cancer Bank, derived from patients with colorectal cancer. Consistent with other sample types, sequencing reached 90% saturation by 80M (150 bp, paired-end) reads for all samples. The resulting datasets show good overall coverage of the genome and excellent consistency for the technical repeats (Supplementary Figure S5). For two of the three samples, we observe high levels of relative enrichment of CpG dense regions of the genome (Supplementary Figure S5A). However, comparison of the three FFPE datasets to similar data for the HT-29 cell line shows good consistency of the profile and the observed ‘background’ of the sequencing dataset (reads lacking CpG sites) is consistently below 5% for all samples. Such an increase in the relative enrichment of regions that are dense in CpG sites is consistent with the expected damage of CpG sites by the FFPE treatment and, feasibly, the protection of these genomic regions (predominantly promoters) from damage in the fixed sample (e.g. by DNA protected within nucleosomal structure).

### Peaks in the Active-Seq profile correlate unmethylated DNA and active chromatin markers

The Active-Seq profile is unique in structure as there are currently no direct comparator technologies that target unmodified DNA for enrichment. Therefore, we benchmarked Active-Seq against established and orthogonal epigenetic methods, namely WGBS and MeDIP-Seq in the NA12878 cell line. As anticipated, the Active-Seq profile shows clear anticorrelation to sites identified as methylated in whole-genome bisulfite sequencing (WGBS) and to those regions of the genome that are enriched in MeDIP-Seq experiments (Figure 2A).

**Figure 2.**
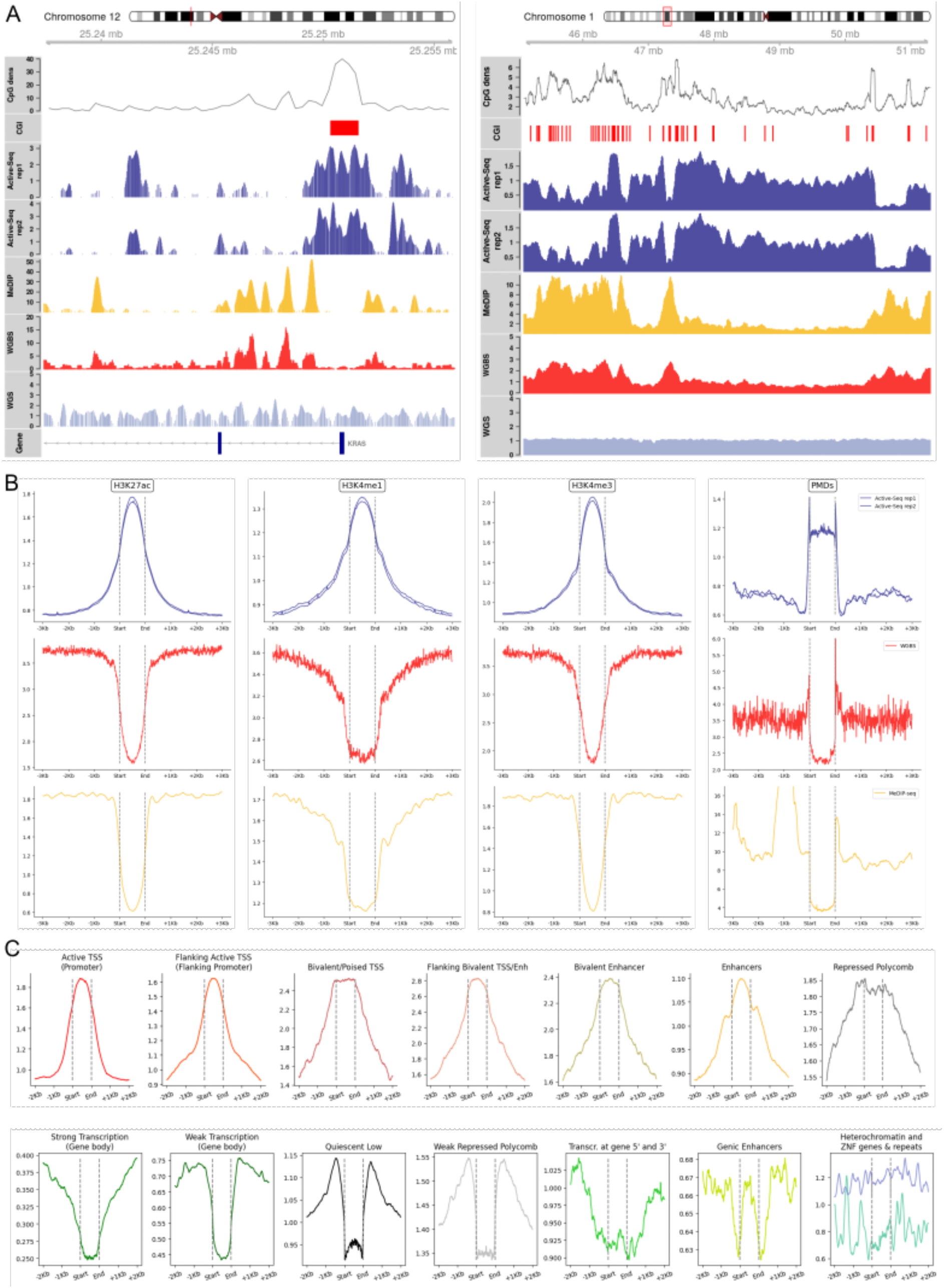
Characterisation of Active-seq signal: **(A)** Genome browser screenshots of Active-Seq coverage (two replicates, blue) compared to MeDIP-Seq (yellow) and WGBS (red) coverage in the KRAS promoter region (left) and in a megabase-scale region of chromosome 1 (right) in NA12878 DNA. **(B)** Profile plot of Active-Seq (blue), MeDIP-seq (yellow) and WGBS (red) at genomic regions associated with histone modification marks (H3K27ac, H3K4me1, H3K4me3) and at partially methylated domains (PMDs) in NA12878 DNA. **(C)** Active-Seq profile plots in genomic features previously identified in the NA12878 genome^*39*^.

Unmethylated DNA typically correlates with active, expressible regions of the genome. Hence, we examined the Active-Seq profile at regions enriched in histone marks for active chromatin. These regions are associated with genomic enhancers (H3K4me1, H3K27ac) and promoters (H3K4me3) and can be aberrantly methylated in diseases^34^. Active-Seq showed a strong enrichment in these regulatory regions; by contrast, MeDIP-Seq shows significant depletion of sequencing reads in these regions (Figure 2B).

Partially methylated domains (PMDs) are prevalent in the cancer methylome and are characterised by wide genomic regions (up to megabases) that bear low levels of DNA methylation^18,19,38^. Active-Seq showed signal enrichment in these regions (Figure 2B), a result which is in line with their intermediate level of methylation and that confirms the ability of this technology to capture the different features that make up the epigenetic profile of a cell type. By contrast, MeDIP-Seq shows poor enrichment for PMDs, which is likely a result of the relatively low CpG site density in these regions in combination with their intermediate methylation levels.

To comprehensively characterize the Active-Seq signal, we analysed the signal profile across 15 chromatin states previously defined in the NA12878 cell line^39^. Active-Seq showed enrichment at transcription start sites (and flanking regions), enhancers, bivalent chromatin as wells as regions repressed by the polycomb repressive complex (Figure 2C). Furthermore, the Active-Seq signal was depleted in regions known to be rich in methylated DNA (intragenic regions of strongly and weakly transcribed genes) and in quiescent/lowly active areas of the genome. Note that, consistent with their highly methylated nature, the Active-Seq profile is depleted in gene bodies that are transcribed (‘Strong Transcription’ in the ChromHMM 15 state model). These genes are characterised by high levels of intragenic DNA methylation, the level of which is enhanced by H3K36me3. This histone modification drives association of the *de novo* methylase DNMT3B to the gene body and it is thought to be a mechanism used to prevent cryptic transcription initiation^40^.

### Active-Seq targets cell-type specific epigenomic markers

Having characterised the features of the Active-Seq signal, we sought to understand its ability to detect differences in profiles from biologically distinct samples. We applied Active-Seq to five cell lines derived from a range of cancers, namely breast (MCF7, HCC1937), colorectal (HT29, SW48) and liver (HepG2). A genome-wide, Spearman correlation analysis of this dataset shows excellent correlation (0.91 – 0.96) of each series of three technical repeats for each of the samples, confirming the robustness of the Active-Seq workflow (Supplementary Figure S6). Each of the cell lines we examined forms a distinct cluster of closely correlated data, with cell lines from similar tissues clustering together (note that HCC1937 and MCF7 cell lines have different subtypes (triple negative, and luminal (ER^+^), respectively)^41^.

To further confirm the ability of Active-Seq to enrich for biologically relevant regions of the genome, we measured the Active-Seq signal at tissue-specific unmethylated markers identified by Loyfer *et al*. in a methylation atlas of healthy human cell types generated using WGBS^12^. These markers (methylation blocks of three or more adjacent unmethylated CpG sites) provide an epigenetic signature for a given cell type since they were identified as being uniquely unmethylated in one cell type versus all other cell types^12^. Figure 3 shows the highly specific Active-Seq signal in the corresponding marker regions identified by Loyfer *et al*. in six different tissue types. Notably, we successfully detected tissue-specific signal despite the heterogeneity of the bulk tissue samples used in our experiments compared to the highly purified, sorted cell types used in Loyfer *et al*. The robustness of these markers combined with the ability of Active-Seq to enrich for unmethylated regions of the genome enabled us to detect a strong, tissue-specific epigenetic signature in these healthy tissues, paving the way to future applications aimed at identifying the tissue of origin of a given sample. Since DNA hypomethylation in enhancers has a known correlation to gene expression levels in cancer^6,42,43^, we anticipate that Active-Seq will be uniquely suited amongst methylation profiling technologies for the study of enhancer methylation and its link to development and disease.

**Figure 3.**
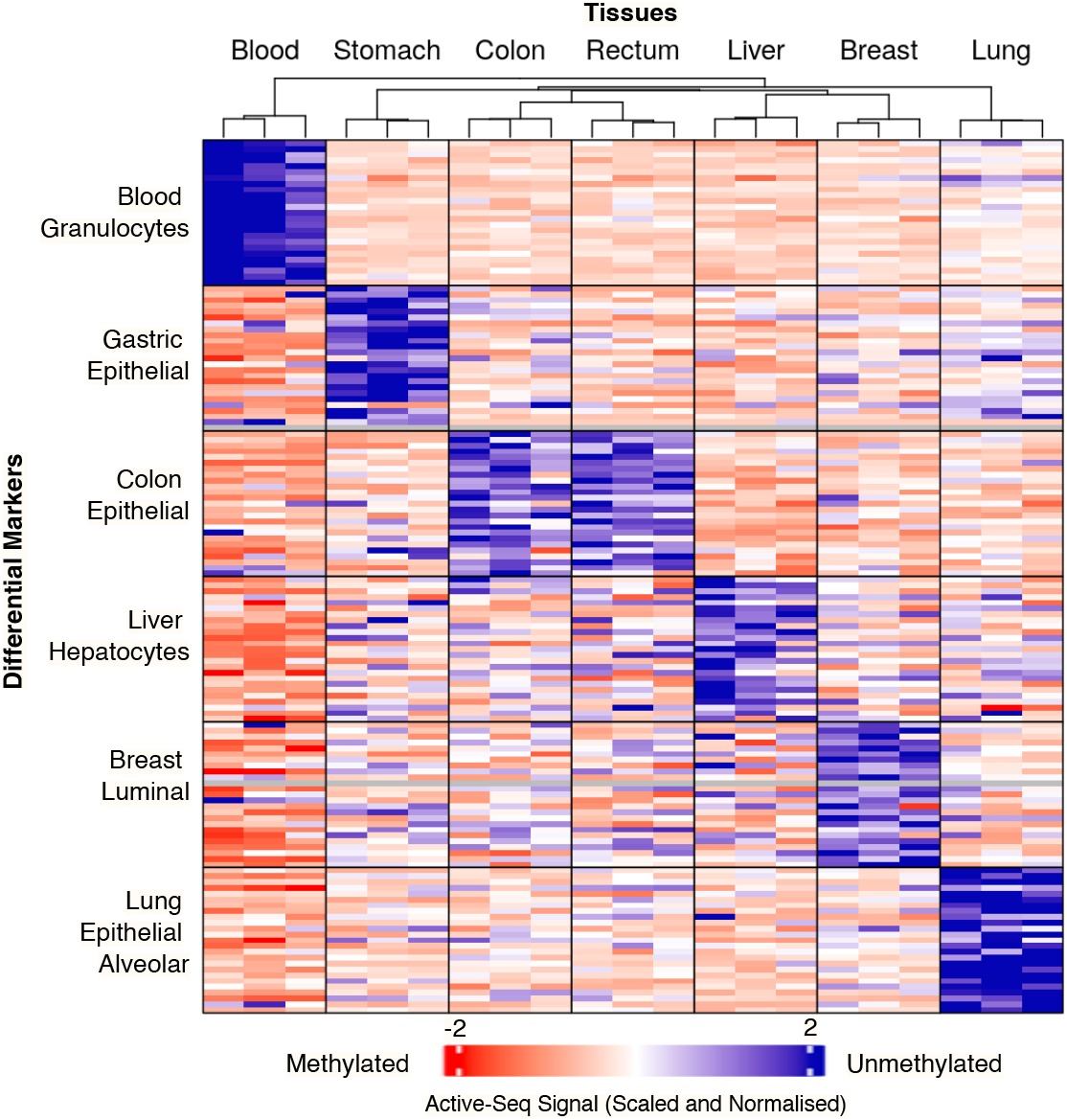
Active-Seq signal at tissue-specific marker regions. Active-Seq enrichment at cell-specific differentially methylated regions (top 25 regions/tissue identified in Loyfer et al). Samples are three technical repeats of Active-Seq except for the blood samples which are derived from the Active-Seq profiles of three individual patients (buffy coat). Tissue-specific differentially methylated regions are grouped according to the tissue/cell type with which they are associated.

### Active-Seq targets genomic regions that are clinically informative

We aimed to establish the potential of Active-Seq as a diagnostic tool to detect changes in methylation status. With this, we aim to extend the application of Active-Seq beyond what was achievable using the technology described by Kriukienė *et al*.^33^, where the variable enzymatic labelling efficiency and uneven representation of the CpG sites of the genome limited such comparative analyses. We undertook a series of experiments using tumour and corresponding normal adjacent tissue from nine patients with colorectal cancer (CRC) and from a collection of six individual patients with six different cancer types. Spearman correlation analysis using the whole Active-Seq profile provides a simple overview for the latter dataset that demonstrates the potentially discriminative ability of Active-Seq for both tissue type and disease status from a patient sample (Supplementary Figure S7).

To highlight the power of Active-Seq in providing insight into CRC biology, we compared the Active-Seq signal generated from tumour and normal adjacent tissues derived from the nine CRC patients. This analysis requires no prior knowledge of a patient’s genomic sequence and makes no assumptions about regions of interest in the genome. We identified differentially methylated regions (DMRs) in cancer versus normal adjacent tissue (see Methods for full details) and identified 4,991 significantly hypermethylated regions and 9,627 significantly hypomethylated regions in this patient cohort (Figure 4A). As anticipated, these hypo- and hypermethylated DMRs cover distinct genomic features. Hypermethylated DMRs highly overlap with transcription start sites (TSS) while hypomethylated regions overlap mainly with intergenic regions and introns (Figure 4B, C).

**Figure 4.**
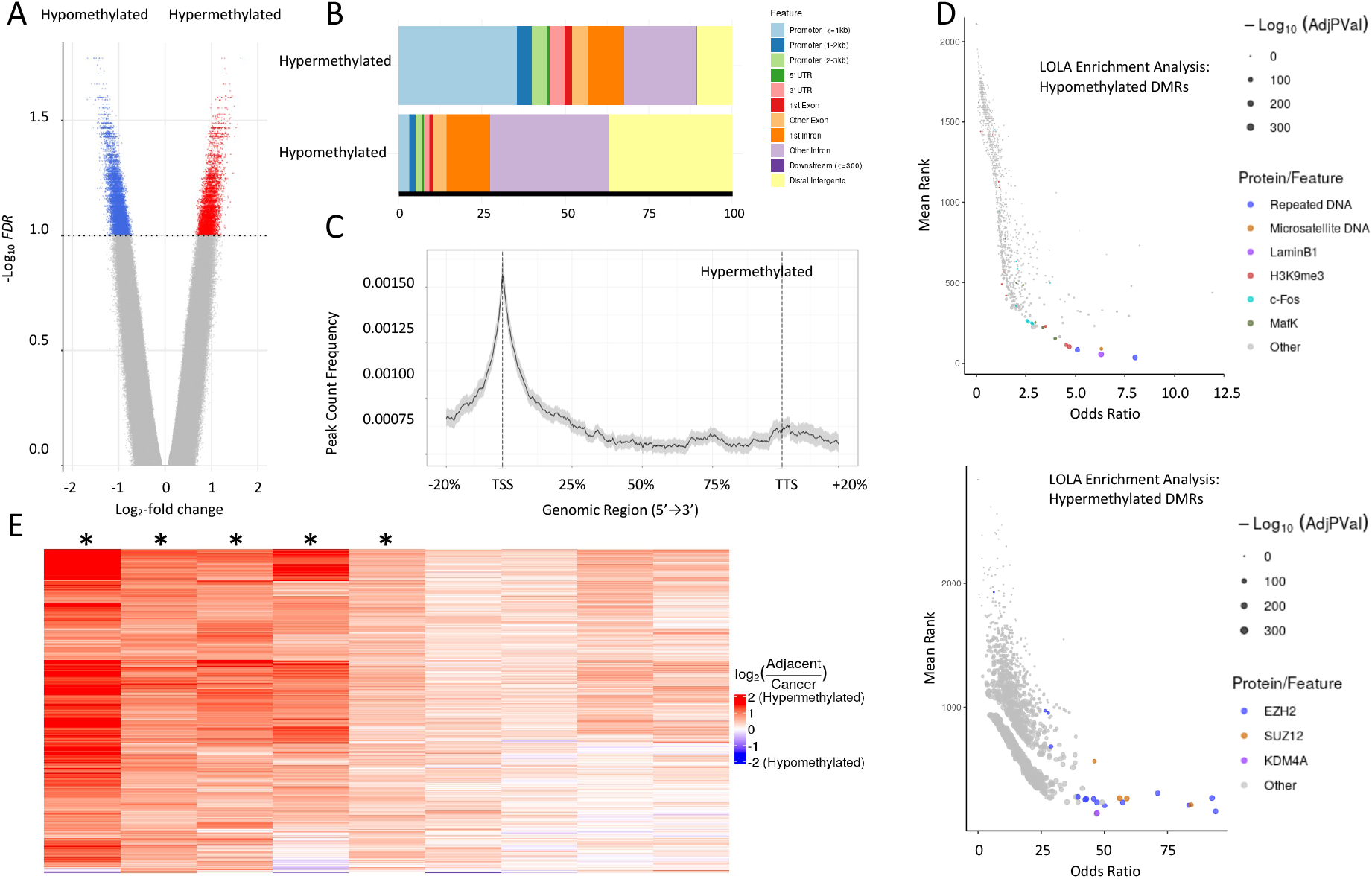
Differentially methylated regions (DMRs) identified in solid tumour samples from nine colorectal cancer patients. **(A)** Volcano plot of hypo-(blue) and hypermethylated (red) DMRs. **(B)** Genomic feature annotation of hypo- and hypermethylated DMRs. **(C)** Frequency profile of hypermethylated DMRs across gene features. **(D)** LOLA enrichment analysis^43^ of hyper-(top) and hypomethylated (bottom) DMRs. **(E)** Hypermethylation level status of the top 500 most variable hypermethylated CpG islands in the nine patient samples profiled with Active-Seq. * = patient identified as potential CIMP^+^ by examining hypermethylation at eight gene promoters^42^.

We then performed a locus overlap analysis (LOLA)^44^ of the hypo- and hypermethylated DMRs with genomic features and transcription factor binding sites (TFBS) associated with this region set. The CRC hypermethylated regions showed a significant overlap with TFBS of the Polycomb repressive complex 2 (PRC2), namely EZH2 and SUZ12, and of KDM4A, an H3K9me3 demethylase. PRC2 is the key enzymatic complex responsible for methylation of the H3K27, a modification associated with gene repression. Targets of the PRC2 complex are predisposed to hypermethylation in adult cancer cells, as a result of these H3K27me3 marks^45–47^.

Mutations to the PRC2 machinery and its mis-regulation have been associated to a spectrum of cancers^48,49^. Indeed, EZH2 is the focus of many attempted therapeutic approaches including the first FDA-approved inhibitor of the PRC2 complex, tazemetostat^50,51^. Similarly, KDM4A is part of the wider KDM lysine demethylase family, and has been widely implicated as a potentially oncogenic enzyme^52^ due to its critical role in the modulation of gene transcription^53^. The Active-Seq hypomethylated DMRs show significant overlap with repetitive DNA regions and lamina-attached domains (as identified by LaminB1 DamID/ChiP-Seq), in agreement with the ability of Active-Seq to detect partially methylated domains (PMDs)^19,3818,1918,19^. The weaker association of these hypomethylated markers (odds ratio 5-10 compared to 25-50 for the hypermethylated markers) is likely a function of the number and the scale of the regions with which the hypomethylated DMRs find overlap.

We further analysed the Active-Seq profiles at the individual patient level. The human genome contains approximately 30,000 CpG islands (CGIs) and about 7.5% of them overlap with the hypermethylated DMRs called in the CRC cohort, while only 0.15% of them overlapped with the hypomethylated ones. Following this observation, we tried to identify patients whose cancers have an epigenetic signature consistent with the CpG Island Methylator Phenotype (CIMP)^54,55^. Colorectal cancer is a complex set of diseases and there have been many different approaches described to stratify patients and predict response to therapy^54,56,57^. Methylation levels at CpG islands, notably in the MLH1 gene promoter, are typically used to identify CIMP positive colorectal cancer patients^58^. We examined promoter hypermethylation at eight genes (*CACNA1G, CDKN2A, CRABP1, IGF2, MLH1, NEUROG1, RUNX3* and *SOCS1)* and made a potential assignment of CIMP^+^ to those patients showing hypermethylation for at least five of these eight gene promoters. This approach tallies with those five patients in our cohort who have a significantly hypermethylated tumour across the most differentially methylated CGIs in the dataset (Figure 4E).

Unbiased and genome-wide profiling of DNA using Active-Seq allowed us to assess both the hypermethylation of promoters-traditionally used to inform patient stratification and prognosis in colorectal cancer^54,55,59^ - as well as the genome-wide hypomethylation, which has been used widely as an early indicator of the disease^21,56,57,60,61^. Active-Seq provides a rich, biologically-relevant perspective on the whole methylome at a fraction of the cost of base conversion chemistries. The enzymatic tagging is a highly efficient process that is non-damaging to the native DNA, leaving the DNA intact for subsequent/ parallel analyses.

### Tumour-informed Active-Seq profiling in liquid biopsy samples

Active-Seq can be applied using an identical workflow on DNA derived from both solid tumour and liquid biopsy samples. As such, it is ideally positioned as a tool for personalised tumour profiling (solid tumour) and subsequent monitoring of disease recurrence or treatment outcome using liquid biopsy, using regions of interest of the epigenome identified in the solid tumour. To test the validity of this hypothesis, we generated profiles from eleven patient plasma samples (seven colorectal cancer and four risk-matched healthy patients) and examined their Active-Seq profiles, identified in solid-tumour samples (at DMRs with an average difference in signal of greater than 1.5-fold). As shown in Figure 5, the two patient cohorts separate clearly using average profiles generated at the hypomethylated DMRs. At hypermethylated DMRs separation of the colorectal cancer and healthy cohorts is less clear, despite the average signal being far greater. Individual DMRs show some promise for resolving the disease status of the two cohorts (Figure 5B) and we will build upon these preliminary results in future and more extensive studies. Critically, Active-Seq facilitates the unbiased identification of epigenetic markers across the genome, without prior selection or knowledge of the patient or their medical condition.

**Figure 5.**
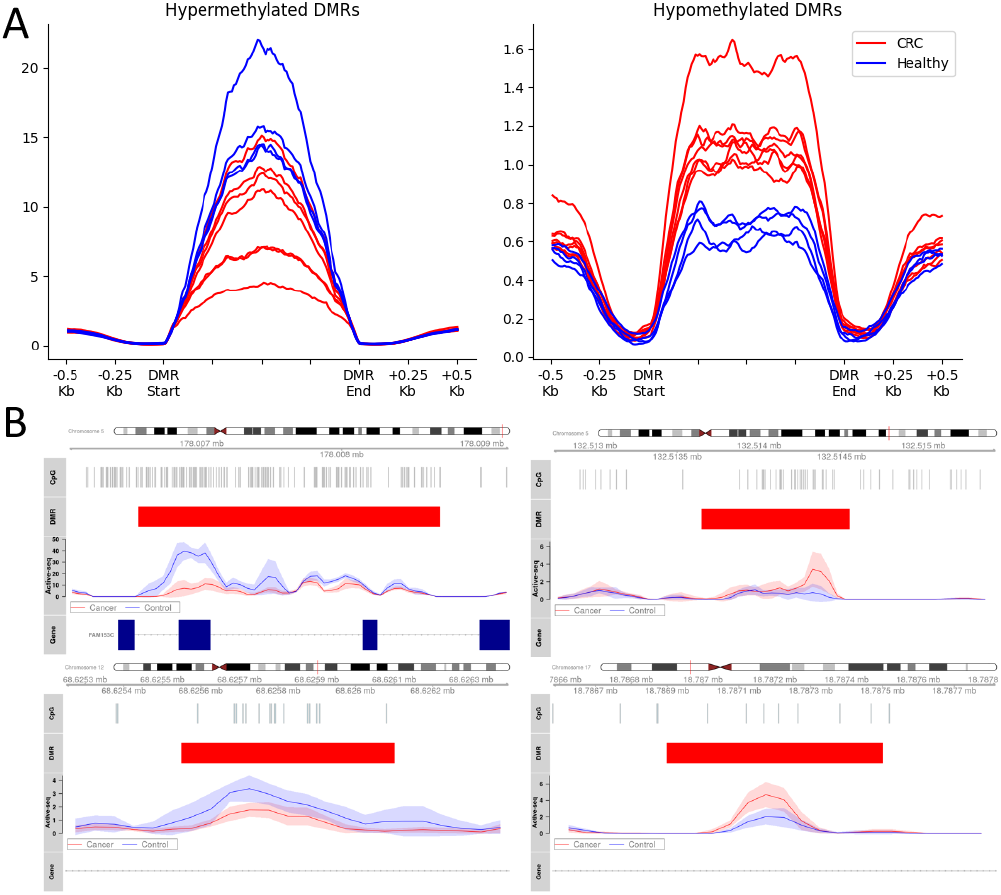
Active-Seq signal in liquid biopsies taken from eleven patients, at differentially methylated regions identified in solid tumour. **(A)** Average signal in patients at hyper- and hypomethylated DMRs for colorectal cancer patients (red) and patients without colorectal cancer (blue). **(B)** Example average profiles across CRC (red solid lines) and healthy (blue) patients for hyper-(left) and hypomethylated (right) DMRs. Standard deviation for the signal is given by the shaded bands.

## Discussion

Active-Seq is an enrichment-based epigenetic profiling technology that, uniquely, focusses on unmethylated CpGs. Technical validation shows that the approach performs robustly across a range of sample types and the simple workflow facilitates reproducible epigenomic profiling with minimal sample loss from input-to-sequencing.

Markers for active chromatin correlate with the regions of unmethylated DNA enriched by Active-Seq. This facet of the technology is unique amongst epigenetic profiling approaches and particularly aligns Active-Seq with recent work highlighting the significance of unmethylated enhancer DNA as a signature of cell biology^12^.

Enhancers are partially (10-60%) methylated in healthy cells and their differential methylation is, in fact, the prevalent perturbation to methylation of DNA at ‘functional’ genomic elements^62–64^. In cancer, hypomethylation of enhancers is more common, being approximately twice as prevalent, in broad range of cancers, as enhancer hypermethylation^62^. Furthermore, the accumulation of enhancer hypomethylation has been shown to correlate with likelihood of metastasis and poor prognosis in lung^65^ breast^66^ and in liver cancer, where hypomethylation of enhancers targeted by CCAAT/enhancer-binding protein-beta correlates with poor outcomes^67^. This is supported by recent single-molecule work that has shown a functional link between DNA methylation status at subset of enhancers and the efficiency of transcription factor binding at these sites^68^. The ability of Active-Seq to target and enrich unmethylated enhancers is a defining aspect of the method and we envisage that this will be critical in its application as a tool to study cancer biology in the future.

Beyond enhancers, the phenomenon of DNA hypomethylation has long been considered a characteristic feature of the cancer genome^20–24^. There is an established link between global and intragenic hypomethylation of the genome, typically measured at LINE-1 elements, which has been associated with poor prognosis in patients with gastric cancers and with advanced colorectal adenomas^56,60,69^ and has been used to monitor the efficacy of DNMT1 inhibitors in clinical trials^70^. Recent works have localised this long-range hypomethylation of DNA to regions known as PMDs, which can be up to megabase-scale domains that correlate with the transcriptionally-repressed genes that predominantly lie within the B-compartment of the genome^71^. Hypomethylation of the PMDs is thought to occur because of their late replication in the cell cycle and is exaggerated in rapidly dividing, cancerous cell populations, as highlighted in a study of over 1500 breast cancer patients^16^. However, as a result of their low CpG density, these sites are poorly enriched in MeDIP-Seq and MBD-Seq and variation in their methylation level, which tends to be of the order of 10-20%, across a broad genomic region, is challenging to measure using base conversion approaches. Active-Seq enables access to these diagnostically useful regions of the genome, which provide an underutilised methylation signature of cancer.

The hypermethylation of specific gene promoters in cancer has been far more extensively studied than the process of genome-wide hypomethylation, that is also a hallmark of cancer^72^. Promoters (linked to H3K4me3 histone mark modification) are stably unmethylated in a healthy cell and are highly enriched in the Active-Seq profile. The abnormal methylation of promoters provides a clear and distinct signature of disease where methylation is typically correlated with gene inactivation. We have identified thousands of significantly hypermethylated regions in tumour derived tissue in the small colorectal cancer study we have performed. Using this signature of hypermethylation, we have been able to identify five patients with highly methylated gene promoters, in the Active-Seq profile, that are consistent with a CIMP^+^ diagnosis for these patients. These initial studies in cell lines and in patient tissue samples demonstrate the potential of Active-Seq as a platform that provides streamlined access to a genome-wide, epigenomic profile. We anticipate that this unique perspective on the methylome will provide new insight for the diagnosis of disease in the near future.

## Conclusion

Active-Seq provides a unique biological perspective on the epigenome and is a robust and straightforward technology that can be broadly applied. The enzymatic approach does not damage DNA nor rely on base conversion. A complete Active-Seq epigenomic profile requires in the region of 15Gbp of sequencing per individual, with no requirement for up-front assumptions about the genomic regions of interest, as is implicit in the use of a panel for genome-wide profiling at the population level using base-conversion technologies. We have shown that the Active-Seq enzymatic chemistry enables unbiased enrichment of unmethylated DNA from a sample for subsequent analysis and that the enzymatic approach is compatible with DNA concentrations that will extend its application toward single-cell analysis (picogram inputs) in the future.

Active-Seq has a unique ability to enrich unmethylated DNA across both the short, highly unmethylated promoter and enhancer regions that are so critically linked to cancer progression and prognosis, as well as the broad, partially-methylated regions of the genome that likely become hypomethylated due to the rapid rates of cell division in tumours. As a result, Active-Seq provides an information-rich, genome-wide profile that we anticipate will enable new understanding of the links between the multitude of processes - from transcription factor binding affinity to genome structural organisation - to which DNA methylation has been linked.

## Acknowledgements

The authors would like to thank Dr Linda Sher and Dr Sang Won Lee, Keck School of Medicine, USC for their kind support and feedback on the manuscript. Biosamples were obtained from the Keck School of Medicine, USC, the Wales Cancer Bank (DOI:http://doi.org/10.5334/ojb.46) and the Northern Care Alliance Research Collection. The Wales Cancer Bank is funded by Health and Care Research Wales. Other investigators may have received specimens from the same subjects.

## Conflict of Interest Statement

LT, CM, AC, AS and JK are employees of Tagomics. JK and RKN are directors for Tagomics. PWL is a consultant and scientific advisor for Tagomics.

## Online Methods

### Shearing of genomic DNA

Where necessary, genomic DNA was sheared to an average of 180 bp in 8 microTube-50 AFA Fiber V2 Strips using a Covaris E220 evolution sonicator.

### DNA tagging reaction

10 ng of input DNA was tagged using the Tagomics Active-Seq reaction kit and according to the manufacturer’s instructions.

### End-repair and A-tailing

The sample was cooled to 10°C and to this, was added 9 μL of End Repair & A-Tailing Master Mix (3.5 μL KAPA End Repair & A-Tailing Buffer, 1.5 μL KAPA End Repair & A-Tailing Enzyme and 4 μL water). The mixture was mixed thoroughly and incubated at 20°C for 30 mins followed by a 65°C incubation for a further 30 mins.

### Adapter ligation

The sample was cooled to 10°C, and to it were added sequencing adapters (2.5 μL of 15μM Stocks) and 22.5 μL of Ligation Master Mix were added (15 μL KAPA Ligation Buffer, 5 μL KAPA DNA Ligase and 2.5 μL water). The mixture was thoroughly mixed and incubated at 20 °C for 30 minutes.

### Capture tag

DNA was mixed with Tagomics’ capture compound and the mixture was incubated at 37 °C for 1 hour with shaking at 500 rpm.

### DNA purification

The DNA was purified from the reaction mixture using AMPure Beads (Beckman Coulter) (washing with 150 μL 80% Ethanol) with final elution in 30 μL PBST (10mM Phosphate Buffer, 1M NaCl, pH 7.5, 0.005 % Tween-20).

### Enrichment of unmodified DNA

Tagged DNA was enriched using 5 μL Dynabeads MyOne Streptavidin C1 beads (ThermoFisher), according to the manufacturer’s instructions. For DNA release, the beads were suspended in 20 μL of Tagomics’ Release Buffer. Released, unmodified (tagged) DNA was transferred to a sterile tube and either stored at -20°C, or used directly for amplification.

### Library amplification

A mixture of KAPA HiFi HotStart ReadyMix (2x, 25 μL), IDT Unique Dual Index Primer Pairs (5 μL of 10 μM Stock) and the enriched DNA library (20 μL) was prepared and subjected to amplification by PCR (up to 13 cycles). The amplified library was purified using AMPure XP Beads and eluted in 10mM Tris-HCl (pH 8.5), 0.01% Tween-20.

### Pooling and sequencing

Libraries were pooled together with 0.1% PhiX and sequenced on an Illumina S4 flow cell on the NovaSeq platform (Source Biosciences).

### qPCR

For each 20 μL qPCR reaction, 1 μL DNA sample, 400nM each of forward/reverse primers specific to the target fragment and 10 μL qPCR mastermix were prepared in triplicate. qPCR was performed on the Azure Cielo 6 thermocycler (Azure Biosystems) with the following conditions: initial denaturation at 98°C for 30 seconds, then 40 cycles at 95°C for 10 seconds and 60°C for 60 seconds with fluorescence detection. Analysis of the acquired fluorescence intensity and subsequent quantification of DNA in the samples was performed using Azure Cielo Manager Analysis Software (V1.0.4).

### Clinical DNA Samples

Cell-free DNA collection was coordinated by the NHS Northern Care Alliance Research Collection, (REC number 18/WA/0368). Blood was collected in K2 EDTA tubes and processed to separate the plasma within four hours of collection. Cell-free DNA was extracted from plasma by Informed Genomics and used directly for Active-Seq. Colorectal cancer samples were provided by Keck Medicine of the University of Southern California. FFPE samples were obtained from the Wales Cancer Bank^73^ which is funded by Health and Care Research Wales. Other investigators may have received specimens from the same subjects. DNA was extracted by YourGene Health and used directly for Active-Seq.

### Processing and alignment of Active-Seq sequencing reads

After sequencing, adaptors were removed from the reads using BBTools^74^ and then aligned to human reference genome HG38 using BWA-MEM2.^75^ Ambiguously aligned reads and those with low mapping scores (MAPQ score < 40) were removed using SamTools^76^. Duplicates were removed with Sambamba (PMID: 25697820) and reads hard-clipped using jvarkit (https://github.com/lindenb/jvarkit). Spearman correlation plots were generated from the processed bam files with deepTools^77^ using a binsize of 1000bp and RPGC normalisation. Saturation figures and CpG density plots were generated for Chr1-22 using the QSEA^78^ and Repitools^79^ R packages. To allow direct comparison of enriched and unenriched samples, Bam files were down sampled to the same sequencing depth using SamTools. High confidence methylated and unmethylated CpG sites used for comparison were taken from the consensus of two whole genome shotgun bisulphite sequencing (WGBS) datasets performed on cell line NA12878 by the same lab (https://www.encodeproject.org/experiments/ENCSR890UQO/), where less than 5% methylation in both datasets was considered to be unmethylated and greater than 95% methylation in both datasets was considered to be methylated.

### Calling of differentially-methylated regions (DMRs)

DMRs were identified using a sliding window approach. The bedtools *makewindows* tool was used to split the hg38 reference genome^80^ in 200 bp windows using a 50 bp step size^81^. Read were summarised in a count matrix using featureCounts, and reads were counted in each window regardless of whether they overlapped multiple windows. Uninteresting windows (as defined as containing less than three sequencing reads per sample on average, or windows that were in the top 0.001% of highest read counts), were filtered out of the matrix using the python package Polars. The filtered matrix was used as input for differential methylation calling using the R package *limma*^82^. Overlapping windows were grouped and correction for multiple testing was performed using the Simes method as implemented in the *csaw* R package^83^. If the FDR was p<0.1 then a region was considered differentially methylated and retained for downstream analysis.

