## Supplementary Information for "Epigenomic profiling of active regulatory elements by enrichment of unmodified CpG dinucleotides"

| | Ladder | Controls | | Cofactor dilution, isomer 1 | | | | Cofactor dilution, isomer 2 | | | | Gel cropped to remove lanes showing activity of M.MpeI with a second, unrelated cofactor | Control -M.MpeI | Control -cofactor | QC lane, enzyme after storage | Control AdoMet + M.MpeI | Ladder |
|---|---|---|---|---|---|---|---|---|---|---|---|---|---|---|---|---|---|
| | | | | 500 µM | 250 µM | 125 µM | 67.5 µM | 500 µM | 250 µM | 125 µM | 67.5 µM | | | | | | |
| R.HpaII | _ | _ | + | + | + | + | + | + | + | + | + | | + | + | + | + | _ |
| M.MpeI | _ | _ | _ | + | + | + | + | + | + | + | + | | _ | + | + | + | _ |
| Cofactor analogue | − | − | − | + | + | + | + | + | + | + | + | | + | _ | + | + | _ |
| Lane | 1 | 2 | 3 | 4 | 5 | 6 | 7 | 8 | 9 | 10 | 11 | | 16 | 17 | 18 | 19 | 20 |

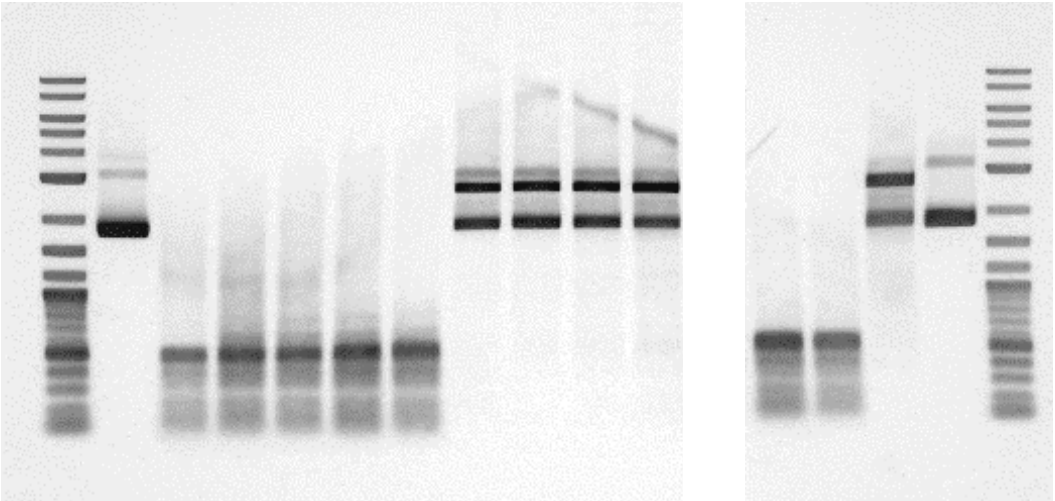

**Figure S1:** Agarose gel showing activity of the M.MpeI enzyme on puC19 plasmid DNA, with the AdoHcyAzide cofactor analogue. DNA is incubated with the methyltransferase and cofactor analogue, purified, then challenged with a restriction enzyme (R.HpaII) which is sensitive to CpG methylation. Lane 2 shows undigested DNA, lane 3 is completely digested DNA. Two cofactor diastereomers are produced in the synthesis and both are tested with M.MpeI here. M.MpeI shows negligible activity with Isomer 1 but near-complete protection of the DNA at cofactor analogue concentrations as low as 68 µM using 'Isomer 2'. Controls show complete digestion of the DNA in the absence of either M.MpeI of the cofactor analogue and complete DNA methylation by M.MpeI in the presence of the native cofactor, *S*-adenosyl-L-methionine.

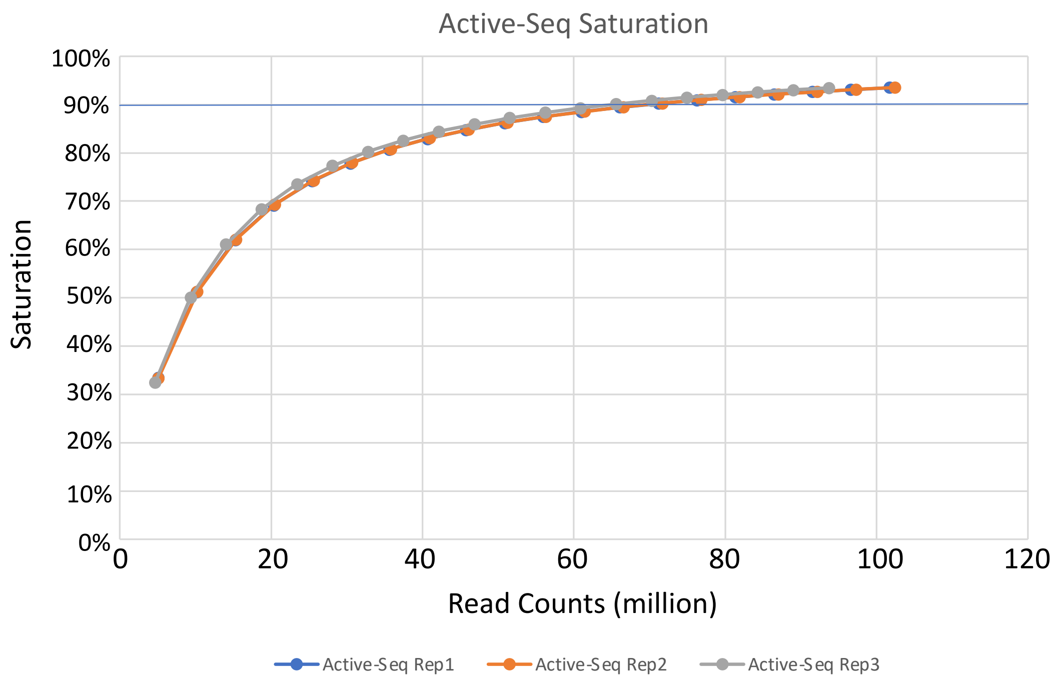


**Figure S2:** Sequencing saturation curves generated for three Active-Seq replicates using the R package MEDIPs[1]. Experiments consistently reach approximately 90% saturation at 70M reads (150 bp paired-end).

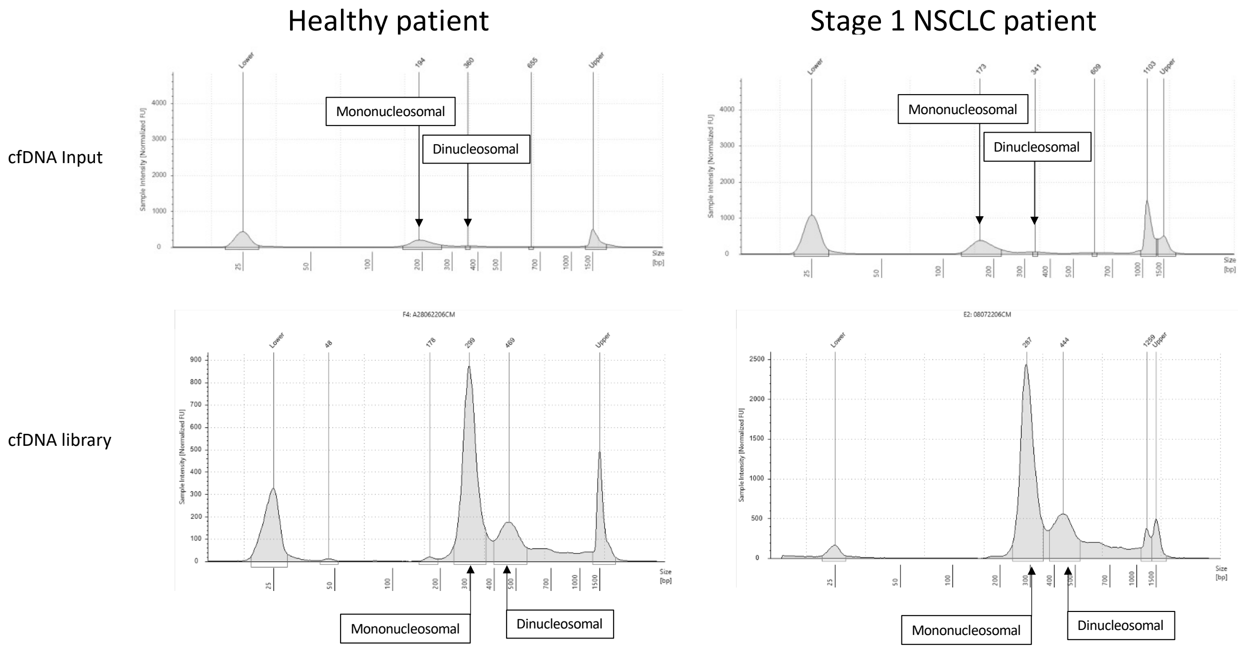


**Figure S3:** DNA Agilent TapeStation (Cell-free DNA ScreenTape) traces showing profiles for the cell free DNA (cfDNA) that was input for Active-Seq profiling (top) and the output from the enriched, amplified libraries (bottom) of the Active-Seq workflow. The mono- and dinucleosomal pattern of the input cfDNA is maintained in the final libraries, the size of which corresponds to the duplicated original strand plus the Illumina P5/P7 adaptors.

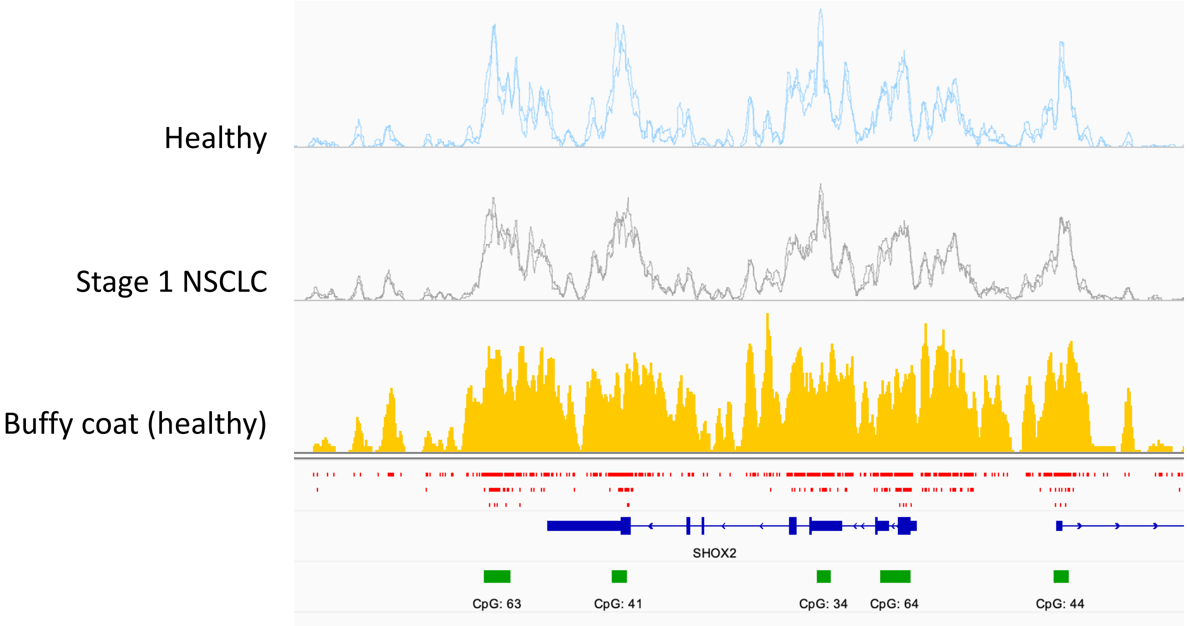


**Figure S4:** Example Active-Seq profiles for a genomic region (SHOX2 gene, a known methylation biomarker for lung cancer) in the healthy (blue) and lung cancer (grey) patients, compared to genomic DNA, extracted from the healthy patient's buffy coat (yellow). Red tick marks show the CpG site density across the gene with CpG islands denoted by green bars. Traces are based on normalised read counts for all profiles and the cfDNA samples are displayed on the same scale for direct comparison.

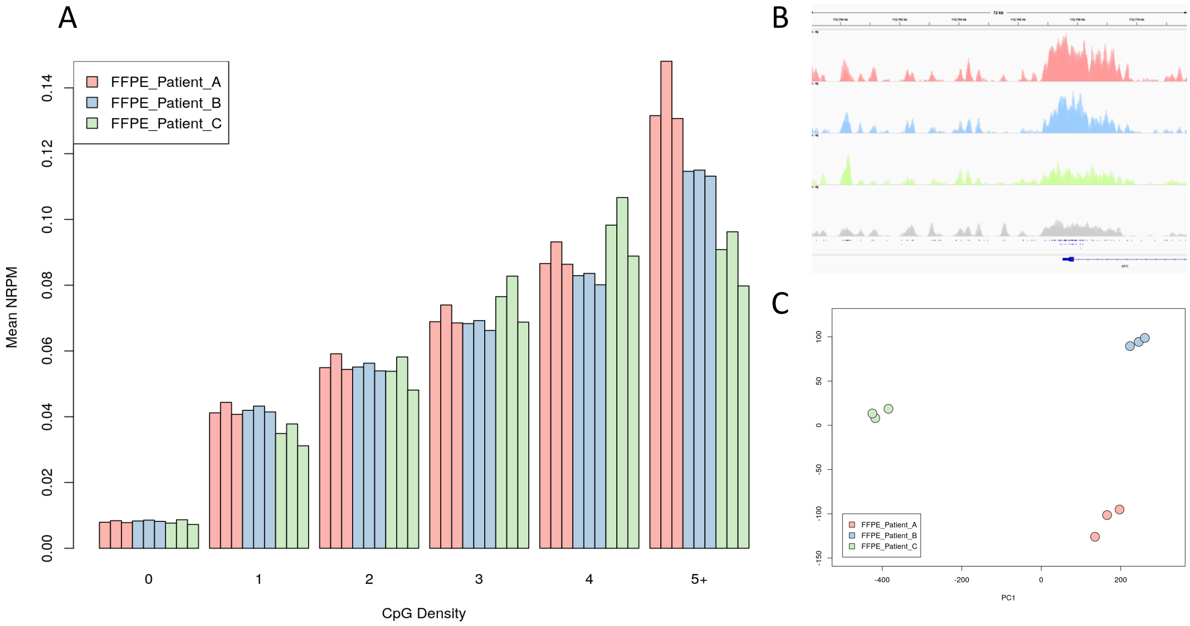


**Figure S5:** Summary of data derived from triplicate repeat experiments using DNA isolated from FFPE. A) Enrichment (normalised read count) as a function of CpG site density (75 bp windows) showing steadily increasing levels of enrichment with increasing CpG density. B) Plots showing normalised read counts across the APC gene transcription start site. Consistent with the enrichment profiles in (A), enrichment of DNA at the CpG-dense gene promoter is more marked for Patient A (pink) than Patient B (blue) than Patient C (green). Profile of the HT-29 cell line (colorectal cancer) shown in grey for comparison. C) PCA plot confirming excellent consistency of the technical replicates of these samples.

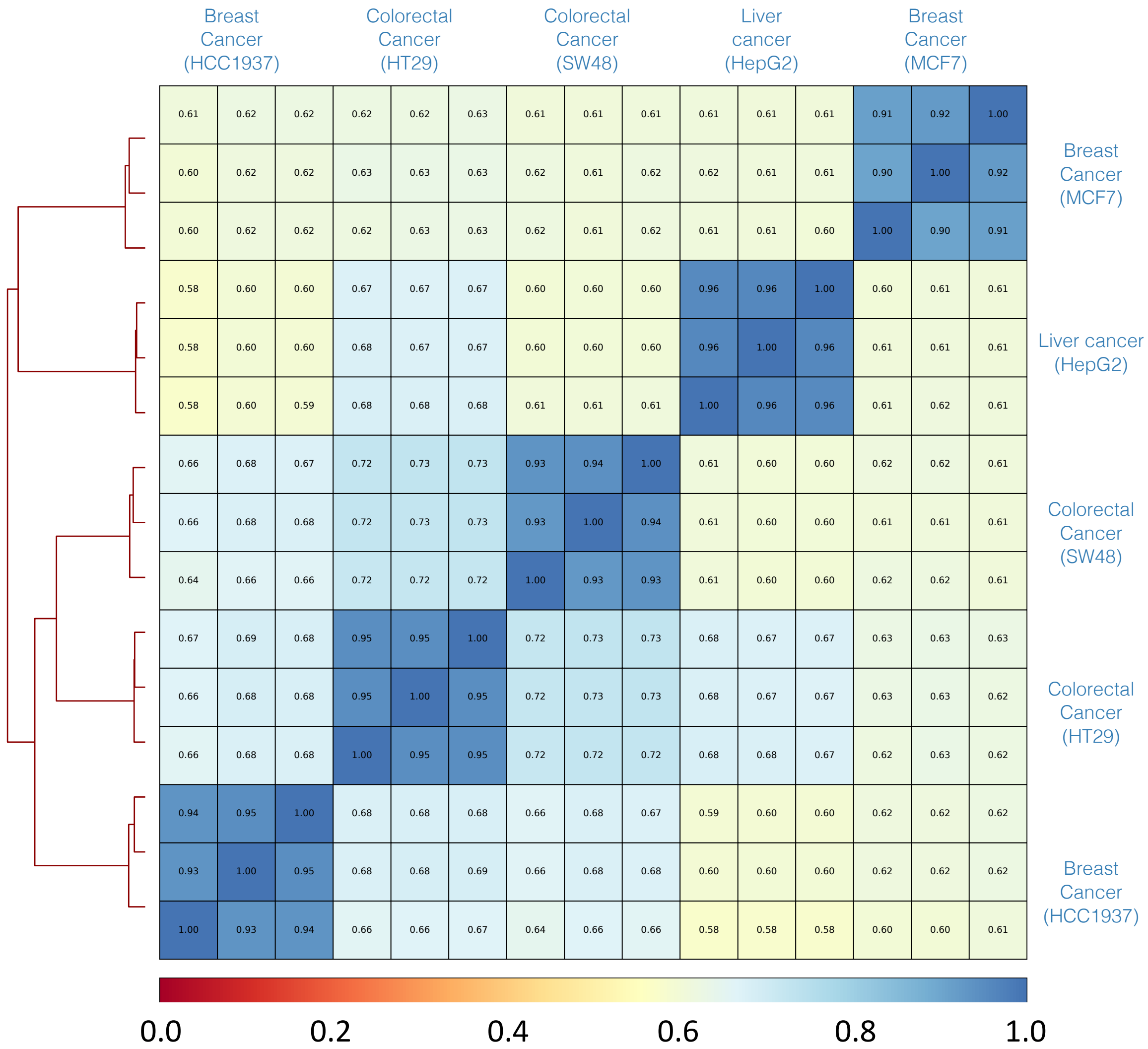


**Figure S6:** Spearman correlation matrix for triplicate repeats of Active-Seq generated from five different cell lines.

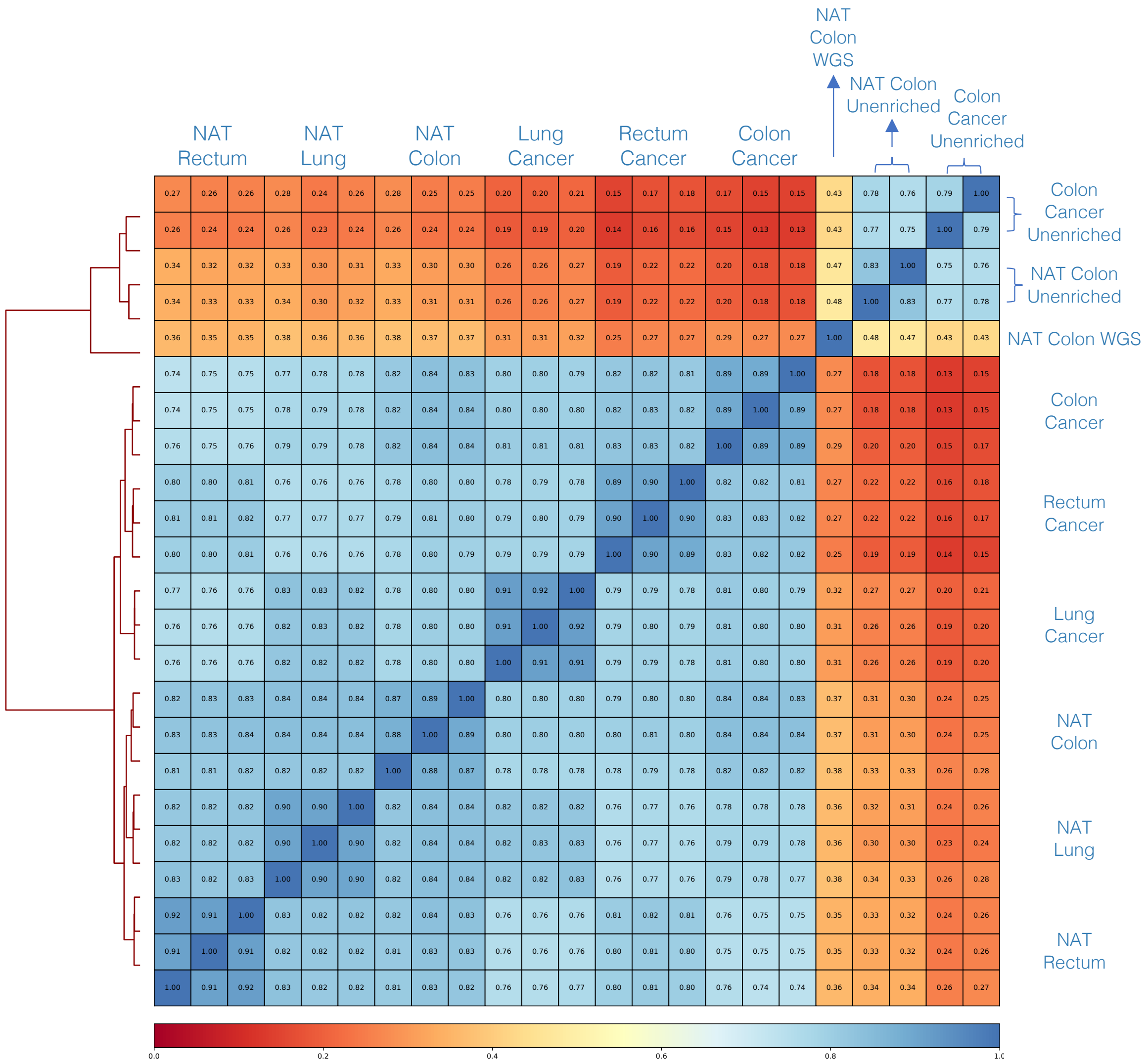


**Figure S7:** Spearman correlation matrix for triplicate repeats of Active-Seq taken from tumour and normal adjacent tissue for patients with colon, rectal and lung cancer. Also shown are the correlations to the unenriched fractions of the genome for colon cancer and to whole genome sequencing for colon cancer.
